# *Pkd1* and *Wnt5a* genetically interact to control lymphatic vascular morphogenesis in mice

**DOI:** 10.1101/2021.03.31.437795

**Authors:** Tevin CY. Chau, Sungmin Baek, Baptiste Coxam, Renae Skoczylas, Maria Rondon-Galeano, Neil I. Bower, Elanor N. Wainwright, Steven SA. Stacker, Helen M. Cooper, Anne K. Lagendijk, Natasha L. Harvey, Mathias François, Benjamin M. Hogan

**Author notes:** Author for correspondence:* Professor Ben Hogan, Organogenesis and Cancer Program, Peter MacCallum Cancer Centre, Melbourne, VIC 3000, Australia.

## Abstract

Lymphatic vascular development is regulated by well-characterised signalling and transcriptional pathways. These pathways regulate lymphatic endothelial cell (LEC) migration, motility, polarity and and morphogenesis. Canonical and non-canonical WNT signalling pathways are known to control LEC polarity and development of lymphatic vessels and valves. *PKD1*, encoding Polycystin-1, is the most commonly mutated gene in polycystic kidney disease but has also been shown to be essential in lymphatic vascular morphogenesis. The mechanism by which *Pkd1* acts during lymphangiogenesis remains unclear. Here we find that loss of non-canonical WNT signalling components *Wnt5a* and *Ryk* phenocopy lymphatic defects seen in *Pkd1* knockout mice. To investigate genetic interaction, we generated *Pkd1*/*Wnt5a* double knockout mice. Loss of *Wnt5a* suppressed phenotypes seen in the lymphatic vasculature of *Pkd1^−/−^* mice and Pkd1 deletion suppressed phenotypes observed in *Wnt5a^−/−^* mice. Thus, we report mutually suppressive roles for *Pkd1* and *Wnt5a,* with developing lymphatic networks restored to a more wild-type state in double mutant mice. This genetic interaction between *Pkd1* and the non-canonical WNT signalling pathway ultimately controls LEC polarity and the morphogenesis of developing vessel networks. Our work suggests that *Pkd1* acts at least in part by regulating non-canonical WNT signalling during the formation of lymphatic vascular networks.

## Introduction

The lymphatic vasculature develops progressively in a process that involves cell specification, cell migration, cell proliferation and morphogenesis (reviewed in Koltowska et al., 2013). In mice, precursor cells along the embryonic veins acquire lymphatic endothelial cell (LEC) fate via the activity of key transcription factors (Francois et al., 2008; Oliver and Srinivasan, 2010; Wigle and Oliver, 1999). The specified LECs sprout out of the veins from approximately 10 days post coitum (dpc) and migrate throughout the embryo to form the lymphatic network (Hägerling et al., 2013; Yang et al., 2012). Early lymphatic sprouting is known to be induced by vascular endothelial growth factor C (VEGFC), signalling through its receptor VEGFR3 (Karkkainen et al., 2004). Subsequent to sprouting, LECs organise, polarise and undergo coordinated cellular and tissue morphogenesis events before forming mature lymphatic vessel networks. These later cellular events controlling vessel morphogenesis and maturation remain to be fully understood.

A growing body of evidence has demonstrated that LEC polarisation and lymphatic vascular morphogenesis can be regulated by WNT signalling components and pathways in mice (Buttler et al., 2013; Cha et al., 2016a; Cha et al., 2018; Lutze et al., 2019; Tatin et al., 2013). The non-canonical WNT/planar cell polarity (PCP) pathway is a highly conserved pathway that coordinates cell and organelle orientation, coordinated cell movements and tissue polarity during development. The PCP pathway is activated by the binding of several WNT ligands (including WNT4, WNT5A, WNT5B, WNT11 and WNT11B), to Frizzled (FZD) receptors and co-receptors; Receptor tyrosine kinase (RYK) and Rar-related orphan receptor (ROR) (Keeble et al., 2006; Lu et al., 2004; reviewed in Roy et al., 2018). Upon activation, various PCP core components, such as Van gogh-like (VANGL) and Cadherin EGF seven-pass G-type receptor (CELSR), are recruited to the activated receptors. These proteins in turn recruit downstream effectors, such as Dishelvelled (DVL), Diversin (ANKRD6), and Prickle (PK), and generate asymmetry to establish cell polarity. Downstream, activation of factors such as RHO kinase or RAC1 influences cell migration and cell motility to co-ordinate tissue morphogenesis (reviewed in Butler and Wallingford, 2017). Canonical WNT signalling occurs when WNT ligands bind to their cognate receptors, initiating the downstream interactions that lead to translocation of B-catenin to the nucleus. This leads to downstream transcription driven by T-cell factor dependent transcription factors (TCFs) (reviewed in Cadigan and Waterman, 2012).

The autosomal dominant polycystic kidney disease (ADPKD) gene *Polycystic kidney disease* (*Pkd1*), encodes the protein Polycystin-1 (PC1), that is mutated in as many as 85% of patients with polycystic kidney disease (Peters and Sandkuijl, 1992). PC1 is a large 11 transmembrane protein with a long N-terminal extracellular region made up of multiple sub-domains and shown to play a role in mechanosensation (Hughes et al., 1995; Nauli et al., 2003; Nims et al., 2003). PC1 localises to the primary cilium, apical membranes and the desmosomes in various cell types and its C-terminal tail binds to Polycystin-2 (PC2 encoded by Pkd2) (Nauli et al., 2003; Yoder et al., 2002). PC2 is a calcium channel that is activated via PC1 function and influences downstream intracellular signalling events (Hanaoka et al., 2000; Nauli et al., 2003; Vassilev et al., 2001). PC1 has been reported to bind directly to core components of the non-canonical WNT/PCP pathway, PAR3 and aPKC, and to regulate cell polarity in kidney epithelia. This role in kidney tubules, regulates the normal intercalation of epithelial cells, tubular extension and morphogenesis (Castelli et al., 2013). PC1 can also influence Ca^2+^ dependent canonical WNT signalling and in *Xenopus* embryos *Pkd1* genetically interacts with *Wnt9a* and *Dishevelled* 2 (Kim et al., 2016).

We and others have found that mice mutant for *Pkd1^−/−^* display defects in lymphatic vascular morphogenesis (Coxam et al., 2014; Outeda et al., 2014). *Pkd1^−/−^* mice display prominent oedema at 14.5 dpc that is typical in animals with lymphatic vascular defects. In the developing dermal lymphatic network, *Pkd1^−/−^* mice exhibit reduced lymphatic vessel branching and dilated/distended lymphatics as a result of the failure of endothelial cells within these vessels to polarise correctly in the direction of vessel sprouting (Coxam et al., 2014; Outeda et al., 2014). Knockdown of *Pkd1* in cultured lymphatic endothelial cells also results in a loss of polarity during cell migration (Outeda et al., 2014). These earlier studies clearly demonstrated an unexpected role for the polycystic kidney disease gene *Pkd1* in lymphangiogenesis and identified it as a potential lymphatic disease gene. However, the precise pathway that *Pkd1* interacts with to control lymphatic vessel development has remained unexplored.

Here, we aimed to better understand the mechanism by which *Pkd1* regulates LEC polarity and lymphatic vascular morphogenesis. We first investigated the endothelial cell-autonomous role of *Pkd1* in valve development, which is known to be controlled by components of the canonical and non-anonical WNT/PCP pathways. We uncovered a requirement for *Pkd1* for normal lymphatic valve development. We then explored the roles of the non-canonical WNT/PCP components *Wnt5a* and *Ryk* in lymphatic development and uncovered phenotypes in knockout mice that were strikingly similar to *Pkd1* mutant phenotypes. Finally, we generated *Pkd1/Wnt5a* double knockout mice and found that these factors genetically interact, being genetic suppressors of each other (a mutually suppressive relationship), during the regulation of LEC polarity, organisation of endothelial cells within developing vessels and in lymphatic vascular morphogenesis. Overall, these genetic studies suggest a role for *Pkd1* in the modulation of non-canonical WNT/PCP pathway signalling in lymphatic vascular development, likely similar to the developmental role of Pkd1 in the morphogenesis of kidney tubules during development.

## Results

### *Pkd1* cell-autonomously regulates lymphangiogenesis and lymphatic valve development

We have previously reported a cell-autonomous role for *Pkd1* in lymphangiogenesis using *Sox18:GFP-Cre-ErT2(GCE)* to generate conditional *Pkd1* knockout mice (Coxam et al., 2014). In the previous study, *Sox18:GFP-Cre-ErT2(GCE);B6.129S4-Pkd1^tm2Ggg^/J* (*Pkd1^f/f^)* mice showed subcutaneous lymphatic network defects but *Tie2:Cre;Pkd1^f/f^* mice did not. To further validate previous findings, we here used *Cdh5:Cre-ErT2,* (which shows activity from 7.5dpc to adult vasculature (Alva et al., 2006; Sörensen et al., 2009)) and generated *Cdh5:CreERT2;Pkd1^f/f^* mice (hereafter referred to as *Pkd1^iECKO^*). During development, Cre activity was induced with serial tamoxifen injections at 9.5dpc, 10.5dpc and 11.5dpc. At 14.5dpc, we examined subcutaneous lymphatics for Prospero homeobox 1 (PROX1), Neuropilin 2 (NRP2) and Endomucin (EMCN) expression. *Pkd1^iECKO^* embryos showed no marked subcutaneous oedema or morphological changes compared with their wild type siblings (**Figure 1A-B**). The subcutaneous blood vessel network was unaffected (**Figure 1C’-D’, K-L**). However, the lymphatic network of *Pkd1^iECKO^* embryos showed reduced complexity with fewer branch points and loops (**Figure 1C-D, G-H**). In the developing dermis, where lymphatics migrate from lateral locations towards the embryonic midline (James et al., 2013), the lymphatic sprouts of the *Pkd1^iECKO^* embryos showed reduced medial migration and the lymphatic vascular front displayed a greater distance to the midline (**Figure 1I**). Furthermore, the dermal lymphatic sprouts were broader and showed a blunt-tipped morphology (**Figure 1E and F**). Combined with our previous work (Coxam et al., 2014), this confirmed that PC1 functions in an endothelial cell-autonomous manner to regulate lymphatic vascular morphogenesis in the developing dermis. At the level of individual cells within developing vessels, the nuclei of LECs in *Pkd1^iECKO^* embryos displayed reduced ellipticity compared with those of control vessels (**Figure 1E, F and J**). Reduced nuclear ellipticity is an indication of cells with reduced motility and a failure to polarise (Hägerling et al., 2013), suggesting a reduction in polarity or motility in *Pkd1^iECKO^* lymphatic endothelial cells.

**Figure 1.**
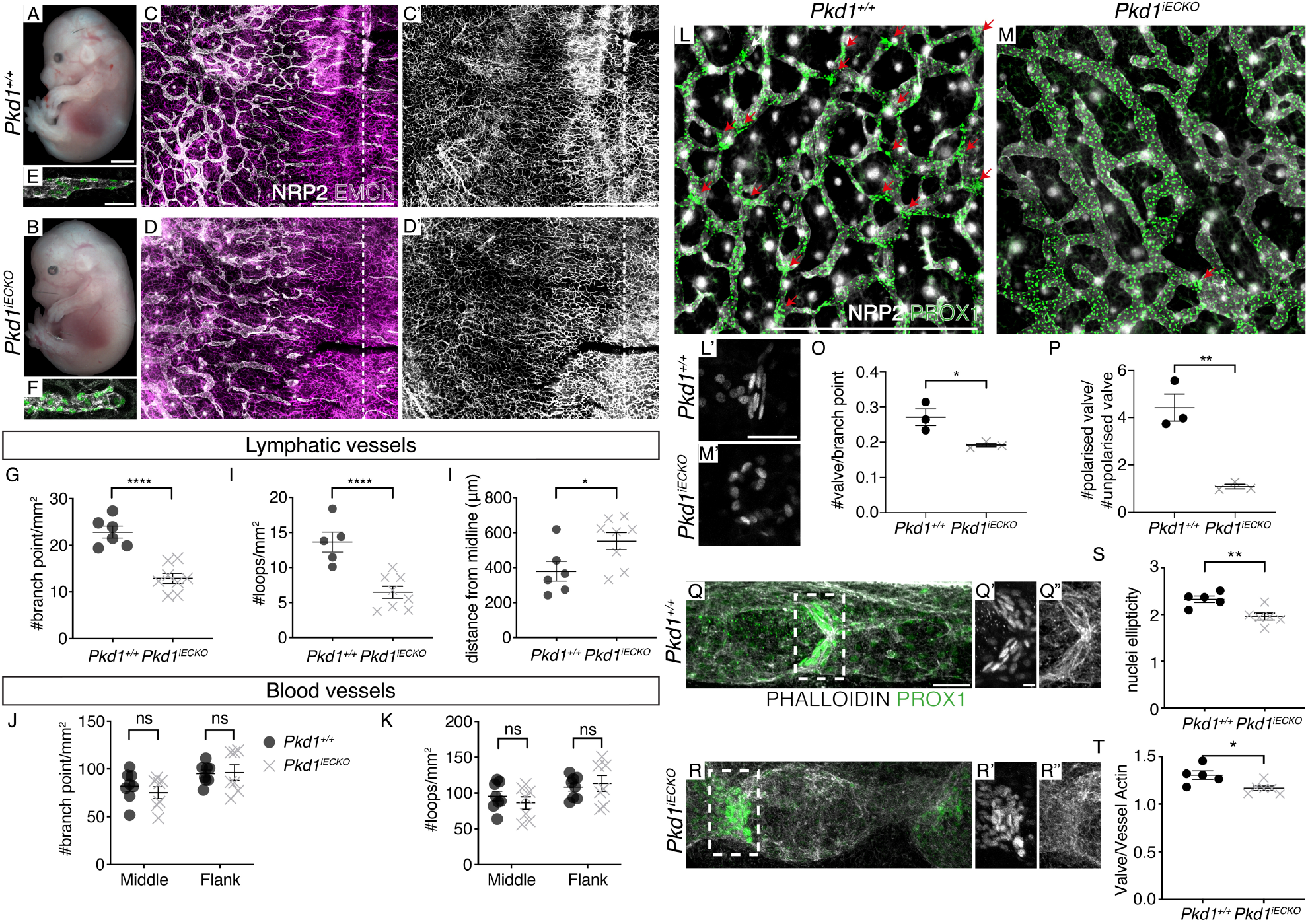
*Pkd1* cell-autonomously regulates subcutaneous lymphatic network and lymphatic valve formation in mice. (A-B) The overall morphology of *Wt* and *Pkd1^iECKO^* mouse embryos at 14.5dpc. Scale bar represents 2mm. (C-D) Subcutaneous lymphatic networks in *Wt* and *Pkd1^iECKO^* embryos at 14.5dpc stained with NRP2 (grey) and EMCN (magenta). Scale bar represents 1000μm. (C’-D’) Subcutaneous blood vessels in *Wt* and *Pkd1^iECKO^* embryos at 14.5dpc stained with EMCN. Scale bar represents 1000μm. (E-F) Subcutaneous lymphatic sprouts in *Wt* and *Pkd1^iECKO^* embryos at 14.5dpc stained with NRP2 (grey) and PROX1 (green). Scale bar represents 50μm. (G-J) Quantification of (G) the number of branch points and (H) loops per mm2, (I) distance of lymphatic sprouts from the midline (μm), and (J) nuclei ellipticity in *Wt* (n=6) and *Pkd1^iECKO^* (n=8) embryos at 14.5dpc. (K-L) Quantification of (K) the number of branch points and (L) loops per mm2 of the subcutaneous blood vessels in *Wt* (n=6) and *Pkd1^iECKO^* (n=8) embryos at 14.5dpc. (M-N) Subcutaneous lymphatic networks in *Wt* and *Pkd1^iECKO^* embryos at 16.5dpc stained with NRP2 (grey) and PROX1 (green). Scale bar represents 1000μm. Arrows indicate lymphatic valves. (M’-N’) Magnification of the mesenteric lymphatic valves in *Wt* and *Pkd1^iECKO^* embryos at 18.5dpc with PROX1 staining. Scale bars represent 50μm. (O-Q) Quantification of (O) the number of valves per branch point, (P) ratio of polarised valves to unpolarised valves, and (Q) nuclei ellipticity of valve LECs in *Wt* (n=3) and *Pkd1^iECKO^* (n=3) embryos subcutaneous lymphatics at 16.5dpc. (R-S) Mesenteric lymphatic valves in *Wt* and *Pkd1^iECKO^* embryos at 18.5dpc stained with Phalloidin (grey) and PROX1 (green). Scale bars represent 50μm. (R’-S”) Magnification of the mesenteric lymphatic valves in *Wt* and *Pkd1^iECKO^* embryos at 18.5dpc with (R’-S’) PROX1 and (R”-S”) Phalloidin staining. Scale bars represent 5μm. (T-U) Quantification for (T) nuclei ellipticity and (U) valve to vessel actin fluorescent intensity ratio in *Wt* (n=5) and *Pkd1^iECKO^* (n=6) embryos at 18.5dpc. Mean±s.e.m. are shown. Student’s *t*-test. ****: p-value<0.0001, ***: p-value<0.001, **: p-value<0.01, *: p-value<0.05

To further study the role of PC1 in LEC polarity, we examined the formation of valves in the dermal lymphatic vasculature. The formation of dermal lymphatic valves requires the local up-regulation of Prox1 in valve-forming territories, the polarisation and the reorientation of developing LECs. In 16.5dpc dermal lymphatics, we observed a marked reduction in valve forming territories in *Pkd1^iECKO^* embryos, as determined by scoring regions of high Prox1 expression in the dermal network relative to the number of network branch points (**Figure 1M-N, O**). We observed that the nuclei of the few lymphatic valves that did form in *Pkd1^iECKO^* dermal lymphatics had reduced ellipticity compared with control embryos, indicative of reduced cell polarity in valve forming territories (**Figure 1M’, N’ and Q**). Quantitatively, there were fewer polarised valves in 16.5dpc *Pkd1^iECKO^* dermal lymphatics than in control embryos (**Figure 1P**) and LEC nuclei failed to reorient perpendicular to the vessel walls.

To examine how PC1 regulates lymphatic valve formation in the mesentery, where valve formation has been better characterised (reviewed in Koltowska et al., 2013; and Kume, 2015; Sabine et al., 2018), we injected tamoxifen at 15.5dpc, 16.5dpc, and 17.5dpc and examined mesenteric lymphatics in 18.5dpc embryos. In wild type siblings, valve LEC nuclei were polarised perpendicular to the lymphatic vessels with actin stress fibres aligned at the lymphatic valves in a stereotypical pattern (**Figure 1R and R”**). The *Pkd1^iECKO^* valve LEC nuclei were identifiable based on high levels of Prox1 expression and formed an initial ring, however the cells within these valve territories failed to orient perpendicular to the lymphatic vessel (**Figure 1S-S”**), and had less elongated nuclei (**Figure 1T**). Actin recruitment or organisation at the lymphatic valves was also reduced in *Pkd1^iECKO^* embryos (**Figure 1R” and S”**), as shown by a reduced ratio of valve-to-vessel actin fluorescent intensity (**Figure 1U**). Thus, PC1 endothelial cell-autonomously regulates the formation of lymphatic valves in the mesentery. The lymphatic valve phenotypes that we observed here and the randomised Golgi orientation in the dermal lymphatic sprouts previously reported (Coxam et al., 2014) strongly suggested a loss of cell polarity in *Pkd1* mutants. The lymphatic valve phenotypes were also similar to valve phenotypes seen in published mouse models such as the *Fat4* conditional knockout mouse or the *Celsr1* mutant mouse, both impacting non-canonical WNT/PCP signalling mesenteric lymphatic valves (Betterman et al., 2020; Tatin et al., 2013).

### The planar cell polarity pathway ligand WNT5A regulates lymphatic endothelial cell polarity during development

The LEC polarity phenotypes observed in *Pkd1^iECKO^* and the previously described interaction between PC1 and PCP components in epithelial cells led us to further investigate the relationship between PC1 and the non-canonical WNT/PCP pathway using genetics. Previous studies have shown that a PCP ligand, WNT5A, is required for dermal lymphatic development (Buttler et al., 2013; Lutze et al., 2019). These studies described lymphatic sprout morphology and structure in *Wnt5a^−/−^* embryos but polarity phenotypes in lymphatic network morphogenesis were not fully described.

We analysed the dermal lymphatics of *Wnt5a^−/−^* embryos at 13.5dpc and observed reduced network formation in the *Wnt5a^−/−^* dermal lymphatics (**Figure 2A-B**). *Wnt5a^−/−^* dermal lymphatics were dilated and the lymphatic sprouts failed to migrate towards the midline of the embryo (**Figure 2C-D**). In *Wnt5a^−/−^* mice, there was an increased number of cells in lymphatic vessels (measured along 150μm of the vessel starting at the leading vessel tips of lymphatic sprouts) and the LEC nuclei displayed reduced ellipticity compared with controls (**Figure 2E-F**). This suggested reduced cell polarity and a lack of LEC migration. To assess whether this phenocopies the *Pkd1*-knockout mice, we investigated LEC polarity more directly. In polarised endothelial cells, the Golgi aligns in the direction of migration, positioning between the nucleus and the direction of migration. Lymphatic vascular mutants with disrupted polarity show a loss of this nuclear-Golgi polarisation in the direction of the embryonic midline (Betterman et al., 2020; Coxam et al., 2014). We examined the polarity of LECs in *Wnt5a^−/−^* embryos by examining Golgi (GOLPH4) localisation relative to the LEC nuclei (PROX1) and the midline (**Figure 2I-J**). The Golgi body aligned in the direction of migration in wild type sprouts, but this was reduced in *Wnt5a^−/−^* mice (**Figure 2K-L**), indicating a loss of cell polarity in *Wnt5a^−/−^* LECs. These phenotypes were in line with the report from Lutze et al. (2019), and further suggest that the loss of PCP signalling, polarity and convergence extension leads to dilated lymphatic vessels and isolated lymphatic cysts in *Wnt5a^−/−^* mice. The blood vessel network also had abnormal endothelial cell clusters along the midline (**Figure 2A’-B’**) but there were no measurable differences in the number of branch points and loops in blood vessel networks at 13.5dpc, a stage when wild type embryos were yet to form a dense blood vessel network spanning the embryonic midline (**Figure 2G-H**). Overall, these phenotypes suggested that WNT5A, a PCP pathway ligand, regulates lymphangiogenesis by controlling cell polarity, displaying a similar but more severe phenotype than *Pkd1^iECKO^* mice.

**Figure 2.**
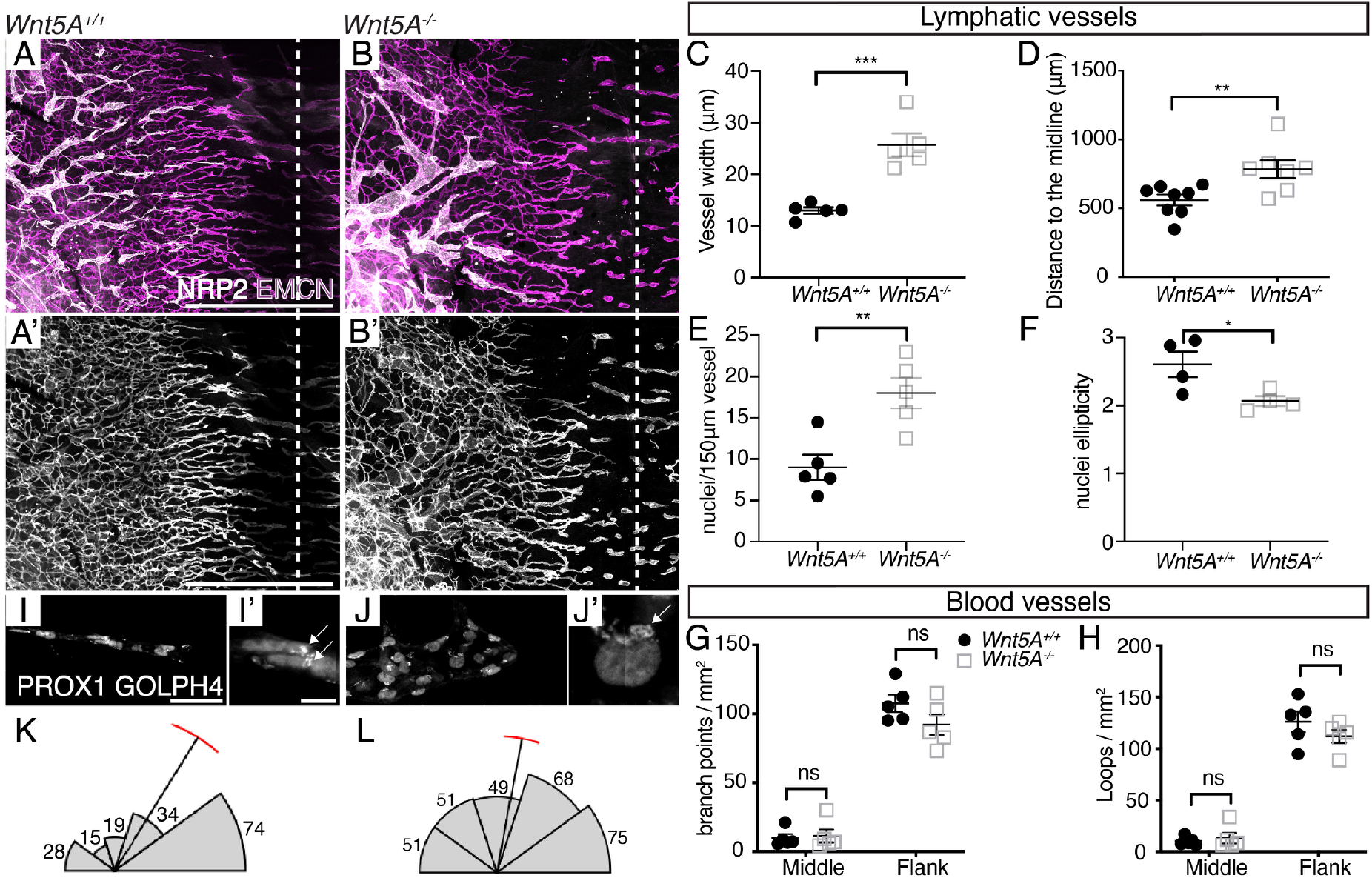
WNT5A mutant mice show subcutaneous lymphatic network and cell polarity defects. (A-B) Subcutaneous lymphatic networks in *Wnt5a^+/+^* and *Wnt5a^−/−^* mouse embryos at 13.5dpc stained with NRP2 (grey) and EMCN (magenta). Scale bar represents 1000μm. (A’-B’) Subcutaneous blood vessel networks in *Wnt5a^+/+^* and *Wnt5a^−/−^* mouse embryos at 13.5dpc stained with EMCN. Scale bar represents 1000μm. (C-F) Quantification of (C) vessel width, (D) distance from midline, (E) the number of nuclei/150μm vessel, and (F) nuclei ellipticity of subcutaneous lymphatic vessel networks in *Wnt5a^+/+^* (n=5) and *Wnt5a^−/−^* (n=5) embryos at 13.5dpc. (G-H) Quantification of (G) the number of branch points and (H) the number of loops of subcutaneous blood vessel networks in *Wnt5a^+/+^* (n=5) and *Wnt5a^−/−^* (n=5) embryos at 13.5dpc. (I-J’) Subcutaneous lymphatic sprouts in *Wnt5a^+/+^* and *Wnt5a^−/−^* mouse embryos at 13.5dpc stained with PROX1 and GOLPH4. Scale bar represents 50μm. (I’-J’) Magnification of a nucleus and its Golgi body in the stained subcutaneous lymphatic sprouts. Scale bar represents 5μm. (K-L) Quantification of Golgi body direction relative to the migration direction of lymphatic sprouts in *Wnt5a^+/+^* (n=5) and *Wnt5a^−/−^* (n=5) mouse embryos at 13.5 dpc. Mean±s.e.m. are shown. Student’s *t*-test. ****: p-value<0.0001, ***: p-value<0.001, **: p-value<0.01, *: p-value<0.05

### The planar cell polarity co-receptor Ryk regulates lymphangiogenesis during development

During PCP signalling, WNT5A binds to receptors, including FZD, ROR and RYK, to initiate downstream cellular processes (reviewed in Butler and Wallingford, 2017). WNT5A enhances RYK and VANGL2 binding to establish cell polarity (Andre et al., 2012). RYK and VANGL2 regulate PCP during mouse cochlea and neural tube development (Macheda et al., 2012). To further understand the role of PCP pathway components in lymphatic development, we assessed the role of the co-receptor RYK in lymphangiogenesis. *Ryk* knockout mice display classical PCP pathway phenotypes that include craniofacial and neural tube defects, and misoriented stereociliary bundles of sensory hair cells in the cochlea (Halford et al., 2000; Macheda et al., 2012). Upon examination of the dermal lymphatics, *Ryk^−/−^* embryos showed similar but milder lymphatic defects compared with *Wnt5a^−/−^* mice. The *Ryk^−/−^* mutant dermal lymphatic network had reduced branching and numbers of vascular loops at 14.5dpc (**Figure 3A-B**), lymphatic sprouts displayed a stereotypical blunt-tipped and widened morphology (**Figure 3C-D, E**) and there were increased numbers of cells in each vessel normalised to vessel length, scored as cell number per 100μm of vessel from the tips of lymphatic sprouts (**Figure 3F**). The blood vessel network was not affected in these mutants (**Figure 3A’-B’, G-H**). Overall, this indicates that RYK, a WNT5A co-receptor, is also a regulator of lymphatic vascular morphogenesis during development.

**Figure 3.**
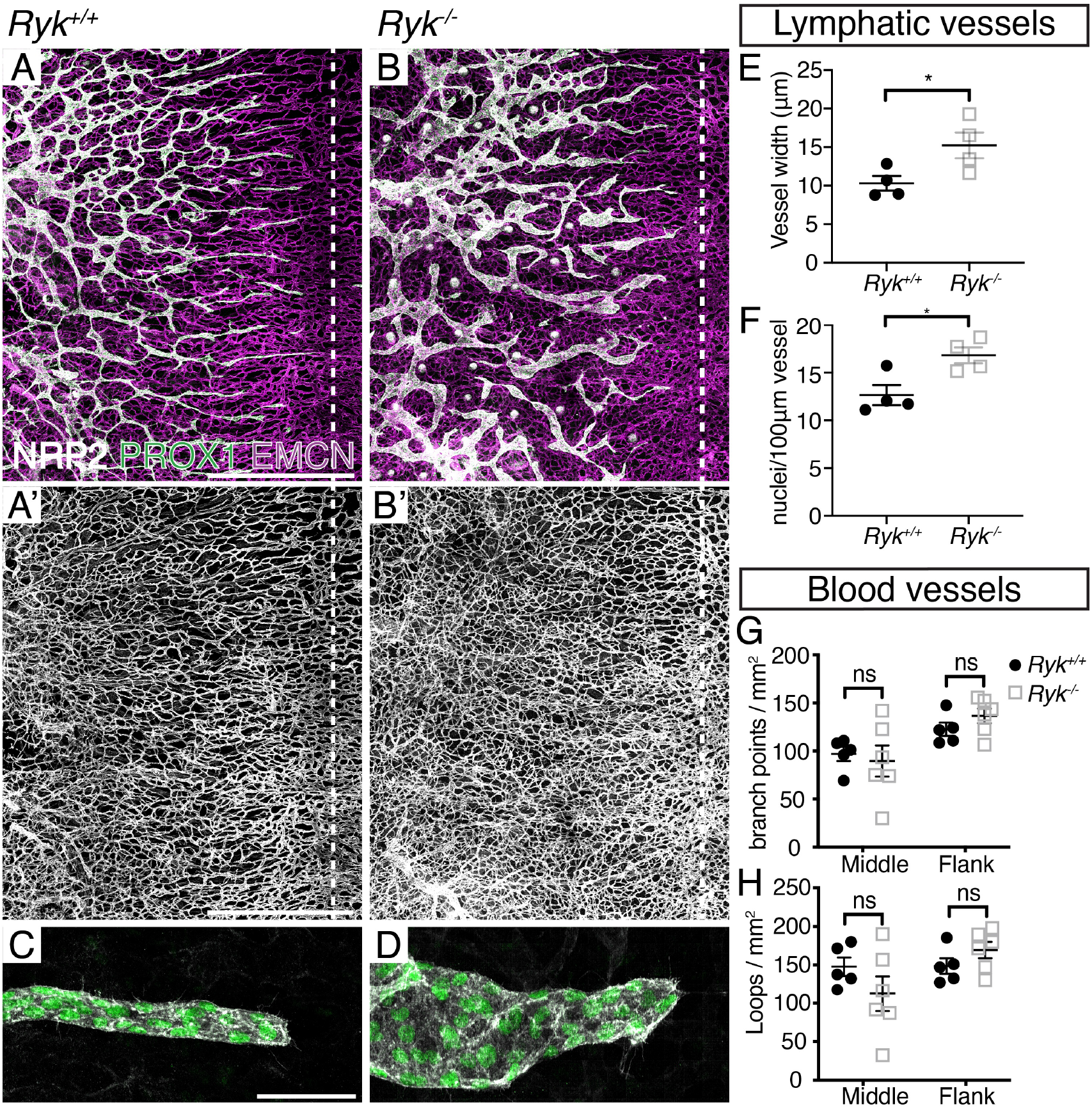
Ryk is required for subcutaneous lymphatic vascular morphogenesis. (A-B) Subcutaneous lymphatic networks in *Ryk^+/+^* and *Ryk^−/−^* mouse embryos at 14.5dpc stained with NRP2 (grey), PROX1 (green) and EMCN (magenta). Scale bar represents 1000μm. (A’-B’) Subcutaneous blood vessel networks in *Ryk^+/+^* and *Ryk^−/−^* mouse embryos at 14.5dpc stained with EMCN. Scale bar represents 1000μm. (C-D) Subcutaneous lymphatic sprouts in *Ryk^+/+^* and *Ryk^−/−^* mouse embryos at 14.5dpc stained with NRP2 (grey) and PROX1 (green). Scale bar represents 50μm. (E-F) Quantification of (E) vessel width and (F) the number of nuclei/100μm vessel of subcutaneous lymphatic vessel networks in *Ryk^+/+^* (n=5) and *Ryk^−/−^* (n=5) mouse embryos at 14.5dpc. (G-H) Quantification of (G) the number of branch points and (H) loops of subcutaneous blood vessel networks in *Ryk^+/+^* (n=5) and *Ryk^−/−^* (n=5) mouse embryos at 14.5dpc. Mean±s.e.m. are shown. Student’s *t*-test. ****: p-value<0.0001, ***: p-value<0.001, **: p-value<0.01, *: p-value<0.05

### *Pkd1* and *Wnt5a* genetically interact to control lymphatic endothelial cell polarity and lymphatic vessel morphogenesis of the dermal vasculature

Given similar phenotypes in the absence of *Pkd1*, *Wnt5a* and *Ryk*, we hypothesised that *Pkd1* may modulate lymphatic vascular morphogenesis by interacting with the non-canonical WNT/PCP signalling pathway. We used genetics to probe for an interaction at the level of vessel phenotype. We generated *Pkd1^+/−^; Wnt5a^+/−^* double heterozygous mice (using a ubiquitous Pkd1-knockout model, not the iECKO model) and in-crossed the double carriers to examine the lymphatic networks for the various genotypes generated. We used a number of quantitative phenotypic measurements to analyse the dermal lymphatics of the 14.5dpc embryos.

As reported previously, both *Pkd1^−/−^* and *Wnt5a^−/−^* lymphatic networks had fewer branch points and loops (**Figure 4A-C**), and the lymphatic vessels were widened (**Figure 4G-I’, M**). We found no evidence that loss of additional copies of *Wnt5a* modified *Pkd1* mutant phenotypes at the level of vessel branch points (**Figure 4B, D-E**). We also found no evidence that loss of additional copies of Pkd1 modified phenotypes at the level of vessel branch points in *Wnt5a* mutants (**Figure 4C, D, F**). However, an examination of lymphatic vessel morphology at the level of vessel width, revealed that developing dermal lymphatics of *Pkd1^−/−^;Wnt5a^+/−^* embryos were thinner than those in *Pkd1^−/−^* knockouts, identifying *Wnt5a* as a dominant suppressor of the *Pkd1* lymphatic vessel width phenotype (**Figure 4J-K, M**). Furthermore, developing dermal lymphatics of *Pkd1^−/−^;Wnt5a^+/−^* double mutants were also thinner than in *Pkd1^−/−^* or *Wnt5a^−/−^* single mutant mice (**Figure 4I-L**). At the level of LECs per vessel (within 100μm from the tips of lymphatic sprouts), *Pkd1^−/−^;Wnt5a^−/−^* embryos displayed fewer cells per vessel than in *Pkd1^−/−^* embryos and were therefore closer to a wild type control vessel (**Figure 4N**). As these data may suggest a genetic interaction at the level of cell polarity and migration, we further quantified nuclear ellipticity in these vessels as a proxy for polarity and motility. *Pkd1^−/−^;Wnt5a^+/−^* and *Pkd1^−/−^;Wnt5a^−/−^* LEC nuclei were more elliptical than those quantified in the *Pkd1^−/−^* LEC nuclei (**Figure 4O**). Finally, we assessed the number of LECs at the tips of leading vessels, as a measure of the blunt tip phenotype. Here we saw not just that loss of alleles of *Wnt5a* suppressed (or partially rescued) the *Pkd1* phenotype, but that loss of a single copy of *Pkd1* dominantly suppressed the *Wnt5a* phenotype as well (**Figure 4P**).

**Figure 4.**
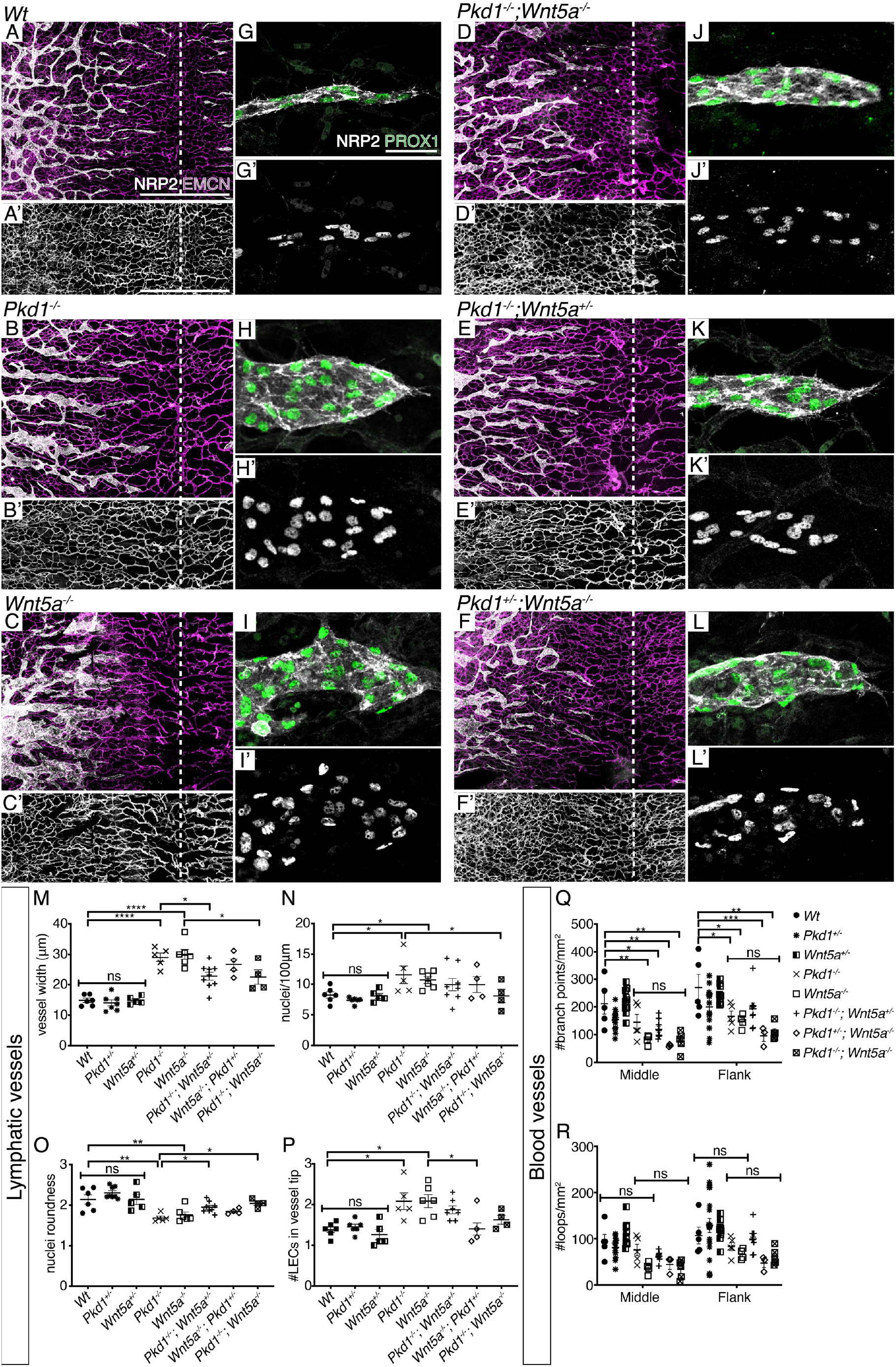
Pkd1 and WNT5A double mutants show less severe lymphatic defects than single mutants. (A-F) Subcutaneous lymphatic networks in (A) *Wt*, (B) *Pkd1^−/−^*, (C) *Wnt5a^−/−^*, (D) *Pkd1^−/−^;Wnt5a^−/−^*, (E) *Pkd1^−/−^;Wnt5a^+/−^*, and (F) *Pkd1^+/−^;Wnt5a^−/−^*, mouse embryos at 14.5dpc stained with NRP2 (grey) and EMCN (magenta). Scale bar represents 1000μm. (A’-F’) Subcutaneous blood vessel networks in *Wt*, *Pkd1^−/−^*, *Wnt5a^−/−^*, *Pkd1^−/−^;Wnt5a^+/−^*, *Pkd1^−/−^;Wnt5a^−/−^*, *Pkd1^+/−^;Wnt5a^−/−^*, *Pkd1^−/−^;Wnt5a^−/−^* mouse embryos at 14.5dpc stained with EMCN. Scale bar represents 1000μm. (G-L) Subcutaneous lymphatic sprouts in (G) *Wt*, (H) *Pkd1^−/−^*, (I) *Wnt5a^−/−^*, (J) *Pkd1^−/−^;Wnt5a^−/−^*, (K) *Pkd1^−/−^;Wnt5a^+/−^* and (L) *Pkd1^+/−^;Wnt5a^−/−^*, mouse embryos at 14.5dpc stained with NRP2 (grey) and PROX1 (green). Scale bar represents 50μm. (G’-L’) Subcutaneous lymphatic sprouts in *Wt*, *Pkd1^−/−^*, *Wnt5a^−/−^*, *Pkd1^−/−^;Wnt5a^−/−^*, *Pkd1^−/−^;Wnt5a^+/−^*, *Pkd1^+/−^;Wnt5a^−/−^*, mouse embryos at 14.5dpc stained with PROX1. (M-O) Quantifications of number of (M) vessel width, (N) the number of nuclei/100μm vessel, (O) nuclei ellipticity of subcutaneous lymphatic networks. (n*Wt*=6, n*Pkd1^+/−^*=8, n*Wnt5a^+/−^*=5, n*Pkd1^−/−^*=5, n*Wnt5a^−/−^*=6, n*Pkd1^−/−^;Wnt5a^+/−^*=8, n*Pkd1^+/−^;Wnt5a^−/−^*=4, n*Pkd1^−/−^;Wnt5a^−/−^*=4) (P-Q) Quantifications of (P) the number of branch points and (Q) loops per mm2 of subcutaneous blood vessel networks. Mean±s.e.m. are shown. One-way ANOVA, Tukey’s multiple comparisons test. ****: p-value<0.0001, ***: p-value<0.001, **: p-value<0.01, *: p-value<0.05

We examined the blood vessel network for similar genetic interaction. The blood vessel network had a reduced number of branch points in all mutant genotypes when compared with wild type embryos in medial and flank regions, likely due to subcutaneous oedema in these mutant embryos. However, there was no change or rescue of this phenotype in the double mutants (**Figure 4A’-C’, Q**). This suggests that the genetic interaction described above is lymphatic restricted and blood vessel phenotypes are likely to be influenced by the sub cutaneous oedema that has been previously reported. We also examined the blood vessel networks between the various genotypes for numbers of loops but found no significant changes compared with wild type (**Figure 4A’-C’, Q**).

Overall, this genetic interaction in lymphatic network morphogenesis between *Pkd1* and *Wnt5a* led to partial phenotypic rescue in various genotypes and provides genetic evidence that *Pkd1* and *Wnt5a* regulate cell polarisation and vessel morphogenesis during lymphangiogenesis by acting in the same genetic pathway.

## Discussion

In this study, we show that *Pkd1*, *Wnt5a* and *Ryk* are important for lymphangiogenesis and LEC polarity. *Pkd1* regulates subcutaneous lymphatic network and valve formation cell-autonomously. The cell migration, elongation and orientation defects reported here and in earlier studies (Coxam et al., 2014; Outeda et al., 2014) suggest that *Pkd1* regulates lymphangiogenesis through controlling LEC polarity. WNT5A and RYK also regulate lymphangiogenesis, with reduced lymphatic network complexity in *Ryk* and *Wnt5a* mutants and loss of polarised Golgi body alignment and reduced nuclei elongation in *Wnt5a* mutants. To further probe the relationship between *Pkd1* and the non-canonical WNT/PCP ligand WNT5A, we generated double mutants and revealed a clear genetic interaction between *Pkd1* and *Wnt5a.* The loss of *Wnt5a* partially rescued (suppressed) the dermal lymphatic phenotypes caused by the loss of *Pkd1*, and in the case of the morphogenesis of vessel tips, vice versa. Overall, these findings report new phenotypes in *Pkd1*, *Wnt5a* and *Ryk* knockout mice and suggest that PC1 acts in the PCP pathway during lymphatic development.

Several studies have suggested the importance of the PCP pathway in lymphangiogenesis. FAT4, a component of the global PCP modules, is required for LEC polarisation and lymphatic development in mice and mutated in the inherited lymphoedema syndrome – Hennekam syndrome (Alders et al., 2014; Betterman et al., 2020). Lymphatic specific deletion of *Fat4* in mice results in lymphatic vessel networks with reduced branch points and increased vessel width. It also leads to reduced mesenteric lymphatic valve formation as does the loss of its ligand Dachsous1 (Betterman et al., 2020; Pujol et al., 2017). LECs in lymphatic specific *Fat4* deletion mice are more rounded, and their Golgi bodies less aligned in the direction of migration and the nuclei (Betterman et al., 2020). Overall, these phenotypes are strikingly similar to those observed in *Pkd1* knockout mice. WNT5A, a ligand of the PCP pathway, is also required for dermal lymphatic morphogenesis (Buttler et al., 2013; Lutze et al., 2019). *Wnt5a*-null mice have reduced numbers of dermal lymphatic capillaries but no difference in LEC proliferation (Buttler et al., 2013). Lymphatic vessels in *Wnt5a*-null mice are cyst-like, non-functional and blood-filled (Lutze et al., 2019). We examined these vessels at 13.5 dpc rather than 14.5 dpc in single mutant animals because of severe morphological defects in these mutants at later stages. These lymphatic vessels show a more severe but still similar phenotype to *Pkd1* knockout mice. In the context of lymphatic valve development, the PCP core components VANGL2 and CELSR1 control valve LEC polarisation and orientation in the mesentery (Tatin et al., 2013). Coupled with our findings above these similar observations support an important role for the non-canonical WNT/PCP pathway in lymphatic network formation and LEC polarity and a role for *Pkd1* interacting with this pathway.

The question remains, how does PC1 interact with the PCP pathway at a mechanistic level? PC1 (encoded by *Pkd1*) can directly interact with PCP components. PC1 complexes with PAR3 to promote PAR3/aPKC complex formation in mouse embryonic fibroblasts (MEFs) and so it is possible the interaction is also direct in LECs (Castelli et al., 2013). However, PC1 also complexes with PC2 to form a Ca^2+^ channel and WNT5A can bind and activate the PC1/PC2 complex to induce Ca^2+^ influx (Kim et al., 2016). It is currently unclear whether WNT/Ca^2+^ signalling contributes to lymphatic development, although it has been shown that shear stress induces Ca^2+^ pulses in LECs (Choi et al., 2017b; Surya et al., 2019) and the Ca^2+^ channel ORAI1 is required for trachea and dermal lymphatic development (Choi et al., 2017a). So it is possible that PC1 may influence Ca2+-dependent WNT signalling, or PCP signalling, or both, to control vessel morphogenesis and the genetic data presented here does not currently distinguish between these possible mechanisms. Of note, canonical WNT signalling is also required for lymphatic development (Cha et al., 2018). The deletion of *β-catenin* and the WNT co-receptors *Lrp5* and *Lrp6* from lymphatics leads lymphatic morphogenesis defects in the dermal vessel network and lymphatic valve formation defects (Cha et al., 2016b; Cha et al., 2018). Thus, there appear to be multiple ways in which WNT signalling cascades can influence lymphatic development and the precise role (or roles) of *Pkd1*/PC1 will require further study in the future.

*Pkd1* is the most commonly mutated gene in autosomal dominant polycystic kidney disease. As such, most emphasis on the role of *Pkd1* in disease has focussed on the kidney. However, this work and our earlier findings suggest that *Pkd1* is a candidate that could influence diseases of the lymphatic vasculature. Of note, in kidney development *Pkd1* also acts by interacting with WNT pathways and the regulation of cell polarity, so there may be common mechanisms at play in both contexts. If PC1 does act via similar mechanisms to FAT4 and DCHS1, given that *Fat4* that is mutated in Hennekam syndrome, it will be interesting to explore the interaction between *Fat4* and *Pkd1* and determine if *Pkd1* may modify disease phenotypes in lymphoedema. It will also be interesting to know if *RYK* or *WNT5A* mutations are associated with primary lymphoedema in humans. Overall, this study confirmed the roles of PC1, WNT5A and RYK in regulating LEC polarity, and identified additional steps of lymphangiogenesis controlled by *Pkd1* (valve formation in the dermis and mesentery). We found that *Pkd1* and *Wnt5a* genetically interact and suggest that *Pkd1* regulates LEC polarity by influencing the PCP pathway. This work thus provides a deeper understanding of the role of *Pkd1* in lymphatic vascular development and points to potential roles in lymphatic disease.

## Materials and methods

### Animals

*Pkd1^f/f^;Cdh5-cre/ERT2* mice were generated by crossing *B6.129S4-Pkd1^tm2Ggg^/J* (*Pkd1^f/f^*; MGI:97603) mice to *Tg(Cdh5-cre/ERT2)1Rha* (MGI:3848982), and breeding carriers to generate subsequent generations and experimental animals. The breeding strategy for the *Pkd1^+/−^* mouse model used in this study was previously described in Coxam et al. (2014). *Wnt5a^−/−^* mice were generated by in-crossing B6;129S7-*Wnt5a^tm1Amc^/J* (mixed background; MGI:98958). *Pkd1^+/−^* and *Wnt5a^+/−^* were crossed to generate double heterozygous carriers. The carriers were incrossed to produce various genotypes of *Pkd1*;*Wnt5a* double mutants and littermate controls.

Timed pregnancies were set up and adult female mice checked for plugs accordingly. To induce Cre recombination and after pregnancy is confirmed, the female is given a dose of 4-Hydroxytamoxifen (Sigma-Aldrich Cat# H6278) dissolved in 90% sunflower oil/10% Ethanol at a concentration of 10mg/mL, during three consecutive days. Dosing of 40mg per kg of body weight was given by intraperitoneal injection. To excise the floxed alleles in *Pkd1^f/f^;Cdh5-cre/ERT* for the dermal lymphatic model, pregnant females were injected at 9.5dpc, 10.5dpc, and 11.5dpc and, for the mesenteric lymphatic model pregnant females were injected at 15.5dpc, 16.5dpc and 17.5dpc.

Genotyping primer sequences: *Pkd1^fl/fl^*-geno Forward-5’-CCGCCTTGCTCTACTTTCC-3’; *Pkd1^fl/fl^*-geno Reverse-5’-AGGGCTTTTCTTGCTGGTCT-3’; *Cre*-geno Forward-5’-CTGACCGTACACCAAAATTTGCCTG-3’; *Cre*-geno Reverse-5’-GATAATCGCGAACATCTTCAGGTTC-3’; *Pkd1*-geno Forward-5’-CCTGCCTTGCTCTACTTTCC-3’; *Pkd1*-geno Reverse-5’-TCGTGTTCCCTTACCAACCCTC-3’; *Wnt5a*-geno Forward-5’-GTTGATTCTGTGTGCCTATTCTGC-3’; *Wnt5a*-geno Reverse1-5’-CCCCGTGGAACTGAGTGTAT-3’; *Wnt5a*-geno Reverse2-5’-TGGA TGTGGAA TGTGTGCGAG-3’. The *Ryk* knock-out mouse breeding and genotyping genotyping was as previously described in Halford et al. (2000).

### Immunofluorescence staining

Embryos were harvested at 14.5dpc or 16.5dpc and fixed in 4% paraformaldehyde in phosphate buffered saline (PBS) at 4°C overnight and then stored in PBS. Dorsal skins from under ear shell to above hind limb were dissected out from the back of embryos and muscle tissue was removed from the skin as previously described by James et al. (2013). The mesenteric membrane was isolated from intestines when harvesting embryos at 18.5dpc. The mesenteric membrane was fixed in 4% paraformaldehyde in PBS at room temperature for 1 hour, and stored in PBS. The skins and mesenteric membranes were blocked in 10% heat-inactivated horse serum in washing buffer containing 100mM maleic acid, 1% dimethyl sulfoxide and 0.1% Triton-X in PBS for 1 hour at room temperature. After blocking, samples were incubated in blocking buffer with primary antibodies overnight at 4°C. Samples were then washed during the day with washing buffer for 5 hours and then incubated in blocking buffer with secondary antibodies. Samples were washed again with washing buffer throughout the day and mounted with Vectashield Antifade Mounting Medium (Vector Laboratories Cat# H-1000-10) on a clear microscope slide. Various combinations of antibodies were used to achieve co-staining of different proteins. Primary antibodies used were: anti-EMCN (Santa Cruz SC-53941), anti-PROX1 (AngioBio 11-002), anti-NRP2 (R&D System AF567). The following secondary antibodies were used: Donkey anti-Goat IgG (H + L) Cross-Adsorbed, Alexa Fluor 488 (Invitrogen, A11055), Goat anti-Rat IgG (H + L) Cross-Adsorbed, Alexa Fluor 647 (Invi-trogen, A21247), Donkey anti-Rabbit IgG (H + L)Highly Cross-Adsorbed, Alexa Fluor 594 (Invitrogen, A21207), Alexa Fluor 488 Phalloidin (Invitrogen, A12379). Stained skins and mesenteric membranes were mounted on glass slides with gelvatol mounting medium and stored at 4°C until imaging.

### Imaging

All mouse embryos were imaged prior to fixation using a Leica M165 FC at 0.73x magnification, illuminated by an external light source. The mounted skins of Figure 1 and 4 were imaged on a Nikon Ti-E Deconvolution Microscope with a 4x Plan Apochromat N.A. 0.2 objective to generate skin scan images for quantifications of lymphatic network formation. The mounted skins and mesenteries of Figure 1 and 4 were imaged on a Zeiss LSM 710 Meta Confocal Scanner with 40x Oil Plan Apochromat N.A. 1.3 and 63x Oil Plan Apochromat N.A. 1.4 objectives to generate vessel images for quantifications all cellular level. The mounted skin of Figure 3 and 4 were imaged with a Zeiss LSM 510 or Zeiss LSM 710 Meta Confocal Scanner with 40x Oil Plan Apochromat N.A. 1.3 and 63x Oil Plan Apochromat N.A. 1.4 objectives.

### Quantification and image analysis

All images were analysed with Fiji (Schindelin et al., 2012). We quantified all branch points and loops of the subcutaneous lymphatic network within 2000μm from the midline in 13.5dpc and 14.5dpc embryos. We randomly selected 5 lymphatic sprouts from each side of the dermal lymphatics, and quantified the ellipticity of 10 nuclei in 14.5dpc embryos in **Figure 1J, 2F and 4O**. We quantified the number of branch points and lymphatic valves of the subcutaneous lymphatic network within 2000μm from the midline in 16.5dpc embryos for **Figure 1O**. We quantified the orientation and nuclei ellipticity (nuclei length-to-width ratio) for the lymphatic valve nuclei that showed upregulated PROX1 (1.5 times of mean fluorescent intensity) for **Figure 1P and Q**. We analysed 5 valves on the primary spokes of mesenteries from each embryo at 18.5dpf that were stained for F-Actin using Phalloidin, lymphatics using PROX1 and veins using EMCN. Since lymphatic valve cells have an upregulation of PROX1 (Srinivasan and Oliver, 2011), all Prox1 upregulated valve nuclei (1.5 times of mean fluorescent intensity) from each valve were quantified for nuclei ellipticity in **Figure 1T**. The intensity of F-Actin and PROX1 staining in **Figure 1U** was measured from the centre Z-section of each Z-stack through each valve based on the location of the nucleus. The ratio of valve to lymphatic vessel mean F-actin staining intensity was calculated. We selected 5 vessels from each side of the skin for the quantification of genetic interaction at a cellular level in **Figure 2–4**. We measured the width of the selected vessels at 10, 25, 50, 75, 100μm from the tip of lymphatic sprouts, and the average width of the 5 measurements was considered as the width of the vessel. The number of LECs in the vessel tip were counted on 5 lymphatic sprouts on each side of the dermal lymphatics in **Figure 4P**. All quantifications were done genotype blinded.

### Statistical analysis

We performed all statistical tests and generated all dot plots using GraphPad Prism 7 except for the rose diagram generated using PAST (Hammer et al., 2001). Specific tests used in each experiment are indicated in the figure legend.

## Acknowledgements

This work was supported by National Health and Medical Research Council of Australia (NHMRC) grants 1079158 and 1146352. BMH was supported by a fellowship (1155221) from the NHMRC. S.A.S. was supported by an NHMRC Program Grant (1053535), an NHMRC Senior Research Fellowship (1154746) and an NHMRC Investigator Grant (1176732) and by funds from the Operational Infrastructure Support Program provided by the Victorian Government, We thank Dr. Kelly Betterman for their help with mesenteric lymphatic staining protocols and Prof. Peter Koopman for providing mice. All images were acquired at the Australian Cancer Research Foundation’s Dynamic Imaging Facility at the Institute for Molecular Biosciences.

## Competing interests

No competing interests declared.

## Funding

NHMRC grants 1079158, 1146352, 1053535. NHMRC fellowships 1155221, 1154746, 1176732.

## Data availability

No publicly available dataset used.

## Notes

### Competing Interest Statement

The authors have declared no competing interest.

